# A generalized structured coalescent for purifying selection without recombination

**DOI:** 10.1101/2024.06.11.598434

**Authors:** Stefan Strütt, Laurent Excoffier, Stephan Peischl

## Abstract

Purifying selection is a critical factor in shaping genetic diversity. Current theoretical models only address scenarios of either very weak or strong selection, leaving a significant gap in our knowledge. The effects of purifying selection on patterns of genomic diversity remain poorly understood when selection against deleterious mutations is weak to moderate, particularly when recombination is limited or absent. In this study, we extend an existing approach, the fitness-class coalescent, to incorporate arbitrary levels of purifying selection. This model offers a comprehensive framework for exploring the influence of purifying selection in a wide range of demographic scenarios. Moreover, our research reveals potential sources of qualitative and quantitative biases in demographic inference, highlighting the significant risk of attributing genetic patterns to past demographic events rather than purifying selection. This work expands our understanding of the complex interplay between selection, drift, and population dynamics, and how purifying selection distorts demographic inference.

---

EVOLUTIONARY inference is essential for unraveling the demographic dynamics that impact genetic variation. Accurate demographic inference requires an appropriate null model; the neutral theory of molecular evolution is a fundamental concept in population genetics, which posits that most genetic variation within species is not caused by natural selection, but by segregating neutral mutations in evolving populations (Kimura and Others 1968). However, purifying selection significantly influences genetic variation, particularly in the absence of recombination (Comeron 2014, 2017; Hartfield 2021). Furthermore, similar to demographic history, purifying selection complicates the inference of positive selection, making both demography and purifying selection essential to understand the evolution of genetic diversity (Stephan 2019). It is thus important to include purifying selection in a realistic model for demographic inference and the detection of positive selection in both non-recombining and recombining regions (Johri *et al*. 2022, 2023).

The evolution of diversity in genomic regions without recombination, such as the human Y chromosome, is constrained by selection acting on their functional sites due to linkage. Uniparental markers in humans have been used to infer sex-specific demography assuming complete neutrality of these chromosomes (Karmin *et al*. 2015), despite evidence of substantial impact of purifying selection on the diversity of the human genome (Pouyet *et al*. 2018). Although the influence of background selection has been noted as a potential confounding factor in the interpretation of differences between human male and female demographic histories, it has been repeatedly rejected as the primary evolutionary force responsible for the observed variations in genetic diversity between human males and females (Lippold *et al*. 2014; Aimé *et al*. 2015; Karmin *et al*. 2015; Poznik *et al*. 2016; Zeng *et al*. 2018). Moreover, it has been shown that purifying selection in the human Y chromosome provides a better explanation of low diversity levels than pure demographic factors and high offspring number variation (Wilson Sayres *et al*. 2014). Therefore, coalescent models that incorporate the effects of purifying selection are crucial for enhancing the accuracy of demographic inference.

Several theoretical approaches have been developed to describe the effects of purifying selection on the inferred demography, but they mainly focus on very weak (O’fallon 2010; O’Fallon *et al*. 2010) or strong selection regimes (Walczak *et al*. 2012; Nicolaisen and Desai 2012;Kaplan *et al*. 1988; Hudson and Kaplan 1994, 1995). Under strong purifying selection, deleterious mutations are efficiently purged, implying a selection-mutation balance which, in turn, enables the derivation of analytical expressions about the equilibrium distribution of deleterious mutations in a population (Haldane 1927; Kimura 1957, 1962; Kimura and Maruyama 1966; Haigh 1978; Otto and Whitlock 2001). Kaplan *et al*. (1988) introduced a structured coalescent model that incorporates the effects of strongly deleterious mutations. Their model of the structured coalescent implements purifying selection by gradually reducing the effective population size in the recent past to a constant smaller size (Hudson and Kaplan 1994, 1995; Walczak *et al*. 2012; Nicolaisen and Desai 2012). If purifying selection is very strong, ad hoc rescaling instead of a gradual reduction well describes the coalescent process.

The equation 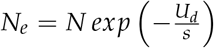 then well approximates the effective population size (Charlesworth *et al*. 1993). Purifying selection is commonly classified as weak or strong, depending on how much it affects the reproductive success of individuals carrying deleterious mutations (Kimura and Others 1968; King and Jukes 1969; Nagylaki 1991). Frequencies of deleterious variants are kept low in populations if selection is strong, but if selection is weak, deleterious mutations can reach high frequencies and even fix in the population. The strength of selection *N s* provides a good measure for selection acting on independent loci, e.g., in scenarios in which recombination is high enough to eliminate linkage and variants with *N s* ≪ 1 do not affect the neutral diversity of a population. However, the absence of recombination complicates the interpretation of the selection strength. If deleterious mutations reach intermediate frequencies, the history of effective population size can be well described by a fitness class coalescent (Walczak *et al*. 2012). A structured coalescent has been derived from the fitness class coalescent to analytically describe the history of effective population size. This model distinguishes between a recent past phase during which the population equilibrates and reaches a more ancient phase where the population is at the aforementioned re-scaled effective population size (Nicolaisen and Desai 2012).

If the deleterious effect of mutations on reproductive success is weak, the relative fitness of individuals may not be significantly affected, such that deleterious mutations are propagated in the population and accumulate over time. A few deleterious mutations eventually fix, i.e., the fittest class in the distribution of deleterious mutations of the population is lost, an event that is referred to as a click of Muller’s ratchet (Muller 1964). A ‘click’ is a direct consequence of purifying selection being too weak to maintain efficient purging, which thus prevents the population from reaching a stationary distribution of deleterious mutations among individuals. Mutation-selection balance is, however, a critical assumption of the structured coalescent, and therefore, it does not apply to scenarios of weak selection due to the fixation of weakly deleterious mutations.

Simulations have shown that the distribution of deleterious mutations can be deterministically described using traveling-wave theory, where the wave maintains a stable profile traveling with a constant velocity through fitness space (Rouzine *et al*. 2003). Although the probability of mutations to eventually fix has been derived for single, independent loci (Haldane 1927; Kimura 1957, 1962; Otto and Whitlock 2001), there is no derivation for multi-locus models without recombination. Numerous approaches have nevertheless been developed to approximate the rate at which the wave travels, corresponding to the click rate of Muller’s ratchet under various conditions, e.g., when selection is very weak and click rates are mainly driven by mutation rates (Haigh 1978; Gessler 1995; Rouzine *et al*. 2008; Metzger and Eule 2013), populations are very small (Lande 1998), or in the presence of recombination (Buffalo and Kern 2024). However, existing theories lack generality in predicting the click rate and the mutation-load distribution under weak purifying selection.

Our primary objective here is to model the coalescent history of populations affected by weak purifying selection in regions without recombination. We are bridging the gap of current theory, which is mainly limited to very weak or strong selection. We extend the structured coalescent based on the traveling wave theory, which allows us to analytically derive the distribution of coalescent times, assuming that both the relative fitness distribution and Muller’s ratchet’s click rate are known. We detail the properties and implications of this model under ideal sampling conditions and describe how purifying selection can generate signatures in the genetic diversity of a population that resemble those of recent bottlenecks.

## Model and Methods

### The structured coalescent for strong purifying selection

The structured coalescent model for strong purifying selection describes the coalescent process of a randomly mixing population by summarizing individuals of the same fitness class into distinct subpopulations (Kaplan *et al*. 1988; Walczak *et al*. 2012). Under mutation-selection balance, an analytical expression of the fitness class distribution exists to describe the frequencies of individuals carrying *k* deleterious mutations (Kimura and Maruyama 1966). Tracing back along genealogies of sampled lineages, either two lineages coalesce, which is only possible if they are in the same fitness class, or a lineage migrates towards the next higher fitness class—the subpopulation of individuals with one fewer deleterious mutation. Lineages can only migrate toward higher fitness classes backward in time because there is no back-mutation (Figure 1). The rate of the deleterious mutations, *U*_*d*_, reflects the migration rate from fitness class *k* to the next less-loaded class *k* − 1 which was analytically derived as *s · k*, with *s* being the deleterious selection coefficient (Nicolaisen and Desai 2012). Since migration occurs on a much faster timescale than coalescence within the fittest class, sampled lineages are unlikely to share a common ancestor except within the fittest class. The separation of these timescales is a requirement to treat the genealogy of sampled lineages as independent. Thus, the independence of lineages is determined by the quantity 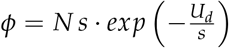 where values of *ϕ* ≪ 1 indicate their independence (Nicolaisen and Desai 2012) validating the assumptions of the structured coalescent model for strong purifying selection.

**Figure 1.**
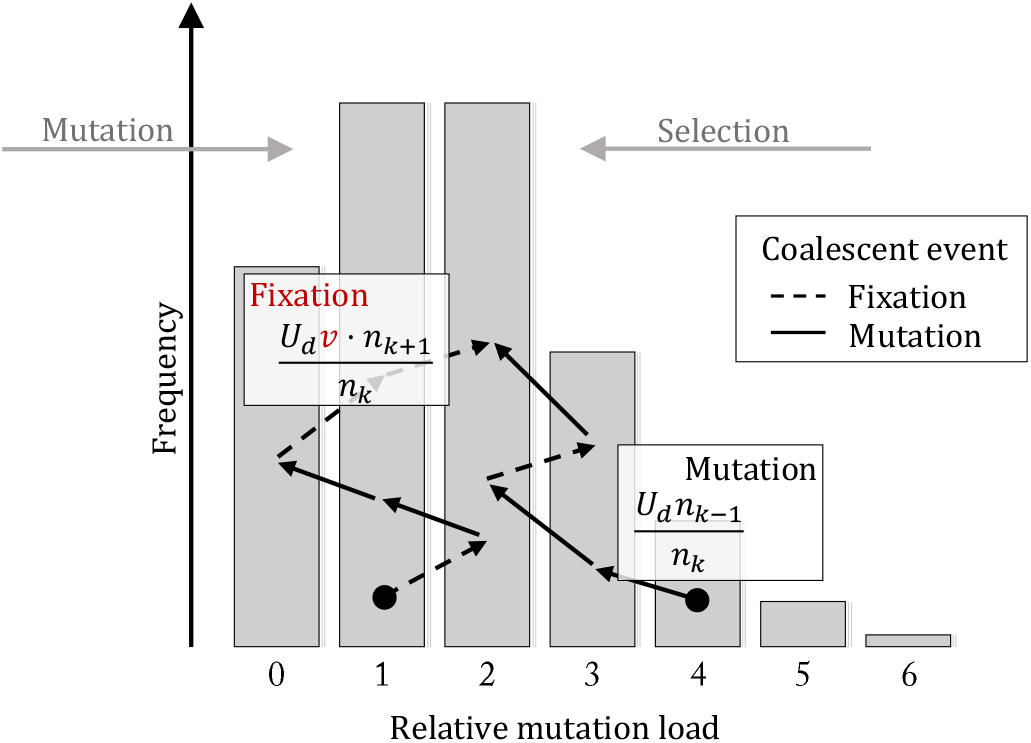
Schematic of the relative mutation-load distribution and its underlying coalescent process in the extended structured coalescent. Deleterious mutations and selection maintain a stationary mutation-load distribution in the population (see Figure 2B). The arrows indicate the coalescent process of two sampled lineages, which backward-in-time migrate through different mutation-load classes until they coalesce. The expected frequencies of sampled lineages being in various mutation load classes change over time. The scaled deleterious mutation rate determines a migration toward less loaded classes (solid arrow), and the scaled fixation rate of deleterious mutations determines migration toward more loaded classes (dashed arrow). Migration rates across classes are a function of the frequencies of the two mutation-load classes between which the migration happens.

### The extended structured coalescent

We modified the structured coalescent into the extended structured coalescent (ESC) to allow for arbitrary strengths of purifying selection (Figure 1). In particular, we consider scenarios where deleterious mutations can eventually fix in the population. Due to the fixation of slightly deleterious mutations, the mutation-load distribution moves toward more loaded classes when evolving forward-in-time (Figure 2A), and the distribution of deleterious mutation frequencies remains stable when expressed relative to the fittest mutation-load class (Figure 2B). Thus, the relative mutation-load distribution 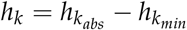 yields a time-homogeneous distribution similar to an evolving fitness distribution under strong purifying selection for which the distribution has been analytically derived by Nicolaisen and Desai (2012) (Figure 2C).

**Figure 2.**
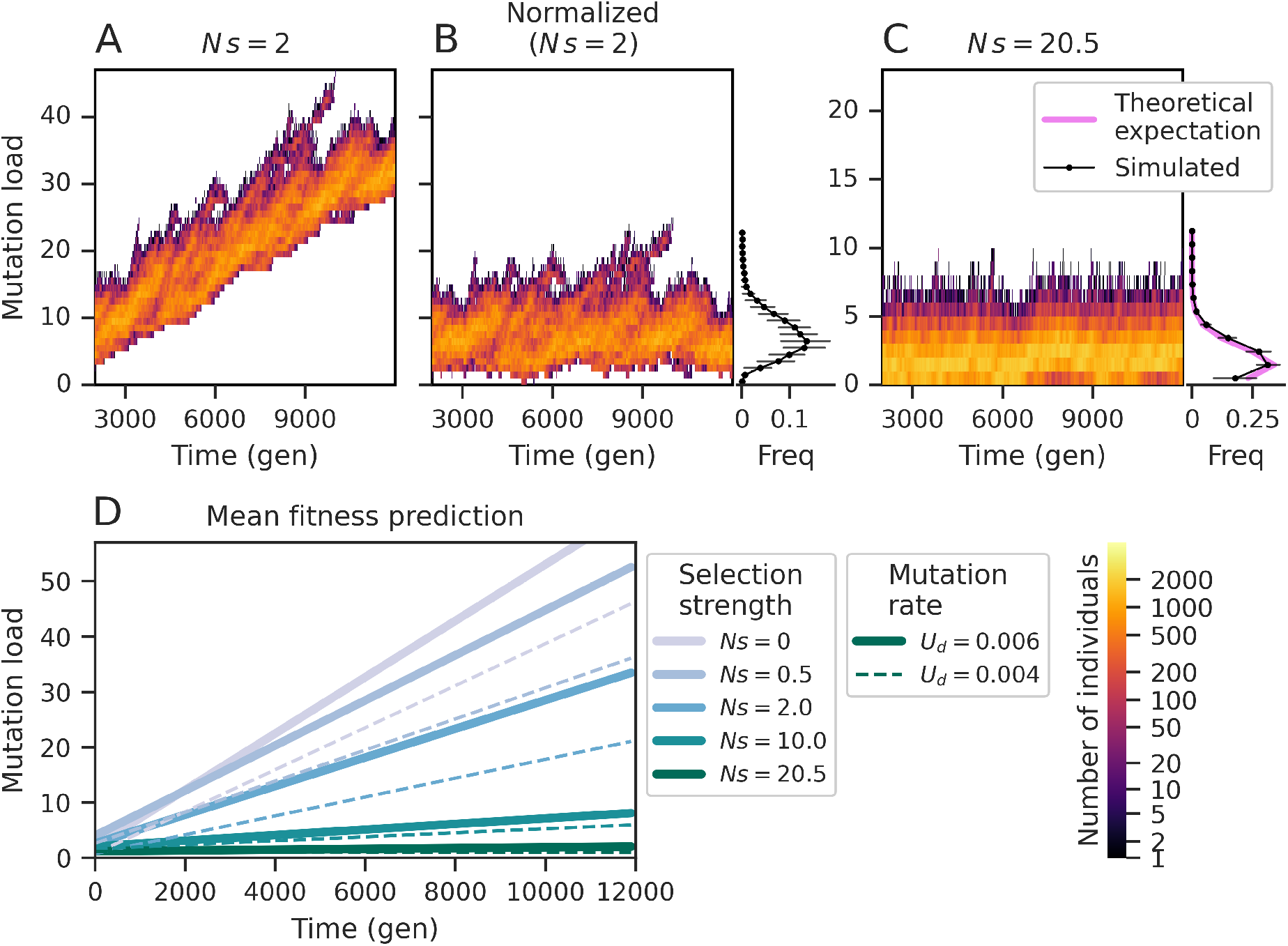
Mutation-load distribution under purifying selection. (A) Traveling mutation-load wave for a population of size *N* = 5000 with purifying selection of strength *s* = 4 *·* 10^*−*4^ and a deleterious mutation rate *U*_*d*_ = 0.006 per generation per locus. (B) Normalization of mutation-load wave shown in panel A resulting in a stationary mutation-load distribution. At each time point, the mutation-load was normalized relative to the predicted mean under a linear model. The global minimum of the mutation-load distribution was redefined as the zero class. The simulated mean number of deleterious mutations of each class is provided in the margin. The error bars show the interquartile ranges. (C) The evolving mutation-load distribution under strong selection parameters for a population of size *N* = 5000 with purifying selection of strength *s* = 4.1 *·* 10^*−*3^ and a deleterious mutation rate *U*_*d*_ = 0.006. The theoretical expectation under the assumption of mutation-selection balance and the simulated mean number of deleterious mutations of each fitness class is provided in the margin. The error bars show the interquartile ranges. (D) The mean number of deleterious mutations is shown for various selection strengths and deleterious mutation rates in an evolving population of size *N* = 5000 under a linear model

Assuming that a relative mutation-load achieves a stationary distribution, we can describe the mutation-load of sampled lineages backward in time by reversing the Markov process of deleterious mutations, similar to the approach proposed for the structured coalescent under strong purifying selection (Nicolaisen and Desai 2012). However, the reverse Markov process requires the addition of a new event that accounts for the fixation of deleterious mutations. Thus, unlike in the structured coalescent, lineages can also move backward-in-time toward more loaded classes (Figure 1, 2A). Therefore, there are two possible migration events to a neighboring mutation-load class, one representing a deleterious mutation event, i.e., a migration towards a less loaded class, and one representing the fixation of a deleterious mutation, i.e., a migration toward a more loaded class, which we can equally describe as a click in Muller’s ratchet. Notably, fixation of a deleterious mutation shifts the relative mutation-load of the classes while the tracked lineage remains unaffected, i.e., former mutation-load class *h*_*k*_ becomes 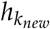 where *k*_*new*_ = *k* − 1 for all *k*. The migration toward more loaded classes is a function of the click rate *v* (*v* for wave velocity) and the stationary frequencies of the source and target relative mutation-load classes. As we cannot obtain the click rate and mutation-load distribution analytically, we estimate the click rate *v* and the stationary mutation-load distribution *h*_*k*_ by simulations (Figure 2D). Following the approach of Nicolaisen and Desai (2012), we then calculate the two types of rates of migration events as 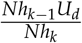 for deleterious mutations and 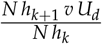 for fixation events, where the relative click rate *v* is obtained by scaling the absolute click rate *v*_*abs*_ to *U*_*d*_. The absolute click rate *v*_*abs*_ is the number of fixed mutations per generation and is scaled to the deleterious mutation rate *U*_*d*_, implying that *U*_*d*_ provides an upper boundary of the absolute click rate, i.e., *v* ∈ [0, 1]. The relative click rate *v* provides an inverse measure of the purging efficacy (Figure 2D, S1), thus providing a numerical value to classify the selection strength regime where *v* = 0 indicates scenarios of strong purifying selection and *v ≈* 1 indicates nearly neutral scenarios.

We obtained the relative click rate *v* and the mutation-load distribution *h*_*k*_ using forward-in-time Wright-Fisher simulations (Figure 2B, C). The deleterious mutation events were modeled as a Poisson point process over time occurring at a rate *U*_*d*_ per individual chromosome per generation assuming an infinite-site model. The selection coefficients of newly-drawn mutations were set to *s*, resulting in an absolute individual’s fitness of 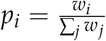, where *k*_*i*_ represents the number of deleterious mutations carried by individual *i*. In each generation, individuals inherit the deleterious mutations from their parents who are randomly chosen from the previous generation with the weighted probability 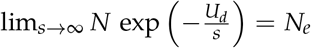 . To obtain *v* from the simulated population, we computed the average change in load per generation, that is the absolute click rate *v*_*abs*_ (Figure 2D). The mutation-load distribution was calculated as the mean of the relative fitness classes *h*_*k*_ (see margins of Figure 2B, C).

The simulated mutation-load distribution is the initial distribution of sampled lineages of the reverted Markov process. We estimated the distribution of lineages in each class of deleterious mutations backward-in-time. We then calculate the expected coalescent rates of each mutation-load class as the probability that two lineages are in the same class divided by that class’s absolute size. The coalescent rates of all classes are summed to provide the effective coalescent rate of the entire population; the inverse of the total coalescent rate estimates the effective population size. Subsequently, the distribution of pairwise coalescent times (*Ψ*_*t*_) is calculated as

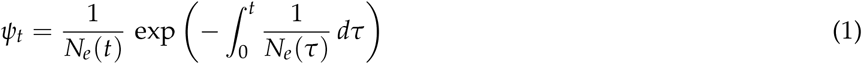

where *N*_*e*_(*t*) is the effective population size of the total population at time *t*.

### Individual-based simulations

To validate our theoretical predictions, we performed forward-in-time Wright-Fisher simulations under purifying selection using the *SLiM 3* software (Haller and Messer 2019). We explored a broad range of parameters, which were chosen to approximate values observed for human uniparental markers and beyond. We determined the parameters reflecting realistic human scenarios as follows: the mutation rate of mtDNA in humans has been estimated lie between 1.3 10^*−*8^ and 1.89 10^−8^ per bp per year, which translates to 3.25 10^−7^ and 4.725 10^−7^ per bp per generation (assuming 25 years per generation) or 0.005 to 0.007 per locus per generation, respectively (Rebolledo-Jaramillo *et al*. 2014; Zaidi *et al*. 2019), by assuming that 90% of 16, 500bp *≈* 15, 000bp are affected by purifying selection (Nicholls and Minczuk 2014). Similarly, for simulating data looking like human Y chromosomes, we used a mutation rate of 1.29 10^−8^ per base pair per generation (Adrion *et al*. 2020) and, following the approach in Wilson Sayres *et al*. (2014), we assumed that 60, 000 sites were non-synonymous (Fujita *et al*. 2010) and obtained an effective deleterious mutation rate of 0.00077 per chromosome per generation. We calculated the coalescent times by drawing 500 pairs of chromosomes from each of 1000 independently simulated populations.

## Results

### The mutation-load distribution under purifying selection

We investigated the properties of the evolving mutation-load distribution by summarizing (1) the fixation rate of deleterious mutations *v*, i.e. Muller’s ratchet’s click rate and (2) the shape of mutation-load distribution *h*_*k*_ (Figure S1, S2, S3). Our simulated results are consistent with theoretical predictions of the mutation-load distribution (see margin of Figure 2C) under strong purifying selection when the click rate is zero (Kimura and Maruyama 1966; Haigh 1978; Hudson and Kaplan 1994). The estimated click rate and the mean number of deleterious mutations depend on three parameters: the selection coefficient *s*, the deleterious mutation rate *U*_*d*_ (Figure S1), and the population size *N* (Figure S2). The click rate decreases monotonously with the selection strength *N s* determining the ‘weak selection regime’ as *v* ∈]0, 1[ (Figure S1). Our click rate estimates are consistent with *ϕ* mapped to the three model parameters *N, s, U*_*d*_. We see several scenarios belonging to the ‘weak selection regime’ (0 < *v* < 1) even if the selection strength *N s* > 10, especially when the population size *N* and the deleterious mutation rates are large (Figure S4). The parameter range belonging to the weak selection regime is restricted to a narrower value range as the deleterious mutation rate *U*_*d*_ and the population size *N* become smaller (Figure S1, S2). The observed relative click rates are consistent with the observed mutation-load distributions: for weaker selection coefficients or higher mutation rates, the relative mutation-load distribution shifts towards more loaded classes, resembling a Poisson distribution with an increasing mean as selection strength decreases (Figure S3). In the case of neutrality (*s* ≈ 0), we expect the mutation-load distribution to follow a Poisson distribution, characterized by a rate of 2 *U T*_*MRCA*_, where *T*_*MRCA*_ denotes the expected time of the most recent common ancestor of the population. Furthermore, we expect that the *T*_*MRCA*_ of a population will be least affected by either very weak or very strong purifying selection when deleterious mutations are nearly neutral or quickly purged, respectively This is because under neutrality the assumption that *N*_*e*_ = *N* is valid and under very strong purifying selection the proportion of individuals in a population being unaffected by deleterious mutations approximates 1 such that 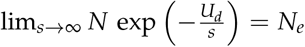 yields the same result as under neutrality (Wakeley 2009; Charlesworth *et al*. 1993).

### Coalescent times and effective population sizes

We calculated the coalescent densities for different scenarios (Equation 1) under our extended structured coalescent and compared the results to estimates from a neutral model, the structured coalescent (Nicolaisen and Desai 2012) and estimates for coalescent times from individual-based simulations (Figure 3). Note that our calculations rely on simulated estimates of the relative click rate and mutation-load distribution (Figure S2, S3). We tested scenarios ranging from nearly neutral, where the standard coalescent is sufficient to describe the coalescent process, to scenarios with strong selection, where the structured coalescent assumptions are met (Figure 3D, E, L). Our predictions of the coalescent density are precise for the entire range of parameters, including neutrality (Figure 3A) and nearly-neutral scenarios (Figure 3H, I). When selection is strong, the extended structured coalescent is equivalent to the structured coalescent since *v* = 0 (Figure 3D, E, L).

**Figure 3.**
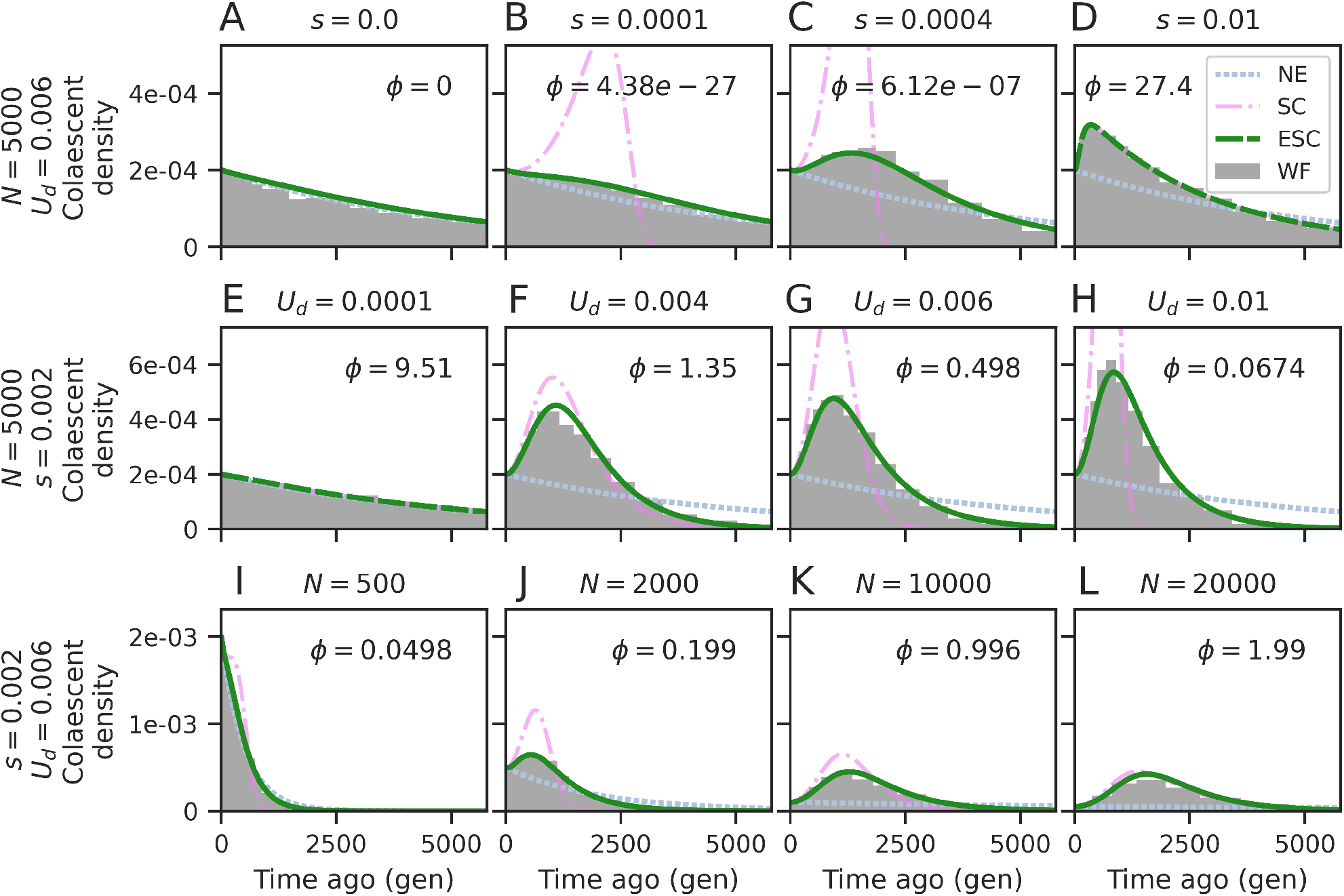
Expected coalescent times under weak purifying selection. Predicted (NE: neutral, SC: structured coalescent, ESC: extended structured coalescent) and simulated (WF: Wright-Fisher forward-in-time simulation using *SLiM 3*) coalescent densities are shown for different sets of parameters. (A-D) Results are shown for fixed parameters *N* and *U*_*d*_ and a varying *s*. (E-H) Results are shown for fixed parameters *N* and *s* and varying *U*_*d*_. (I-L) Results are shown for fixed parameters *s* and *U*_*d*_ and varying *N*. Values of *ϕ* are given for each scenario. The corresponding mutation-load distributions and click rates *v* are provided in Figure S5. The WF coalescent densities are shown for 1000 sampled pairs in 500 independent simulations per condition.

We next show how coalescent densities translate into effective population size histories (Figure 4). We see that the original structured coalescent tends to produce scenarios suggesting a strong recent population expansion, whereas our new extension suggests the occurrence of milder population size expansions. The relative click rate estimated under each scenario provides an accurate indication of the extent to which the effective population size is similar to predictions under strong selection versus neutrality (Figure S5), where values of the click rate *v* tending toward *v* = 0 indicate strong selection and *v* = 1 indicates nearly-neutral scenarios. Our estimate of the coalescent density under neutrality appears very accurate (Figure 3A), however, we observe a very slight recent population expansion (Figure 4A).

**Figure 4.**
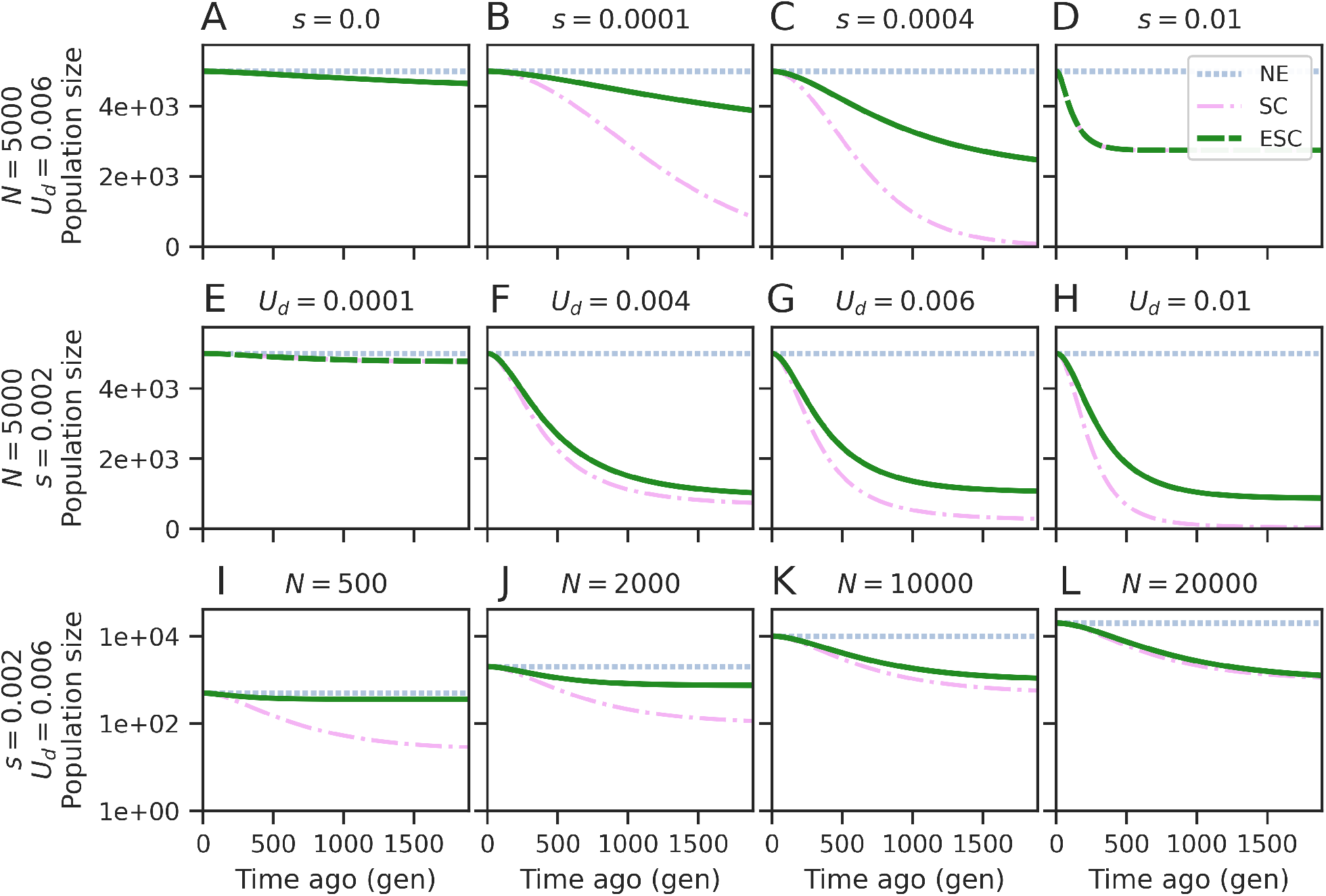
Expected effective population sizes under weak purifying selection. Predicted effective population sizes under different sets of parameters are shown (NE: neutral, SC: structured coalescent, ESC: extended fitness coalescent). This figure complements the coalescent densities in Figure 3 and provides the corresponding history of effective population size for the same sets of parameters, as indicated in the figure panels

### Biases due to sampling

We next investigated the effects of potential biases arising from sampling schemes. We expect that samples from low-frequency classes at the edges of the mutation-load distribution resemble rare haplotypes which may thus appear as genetic outliers and be excluded from analyses. We thus investigated whether sampling lineages from the *n* most frequent mutation-load classes affects the coalescent densities and inferred effective population size histories (Figure 5). To achieve this, we set the initial frequency of the rare classes to zero in the Markov model. This ascertainment introduces a considerable bias in the coalescent times and recent demographic history compared to neutral expectations. Recent coalescent densities are significantly overestimated, particularly when sampling from a single mutation-load class (*n*_*k*_ = 1), as all sampled lineages belong to the same subpopulation in the extended structured coalescent allowing for a significantly quicker coalescence. Consequently, the recent effective population sizes are significantly underestimated before gradually converging towards an unbiased estimate, which would be taken as evidence of a recent population crash for *n*_*k*_ = 1 or a recent transient population size increase for *n*_*k*_ = 5 or *n*_*k*_ = 10 (Figure 5D, E). This biased inference is most prominent when sampling from a single mutation-load class but becomes almost negligible when sampling from ten or more classes since the sampled lineage distribution per class better approximates the relative distribution of deleterious mutations for larger samples. Under stronger selection, the total number of mutation-load classes in the population is relatively small, implying that sampling from a small number of mutation-load classes may suffice to perform accurate inference (Figure 5C, F).

**Figure 5.**
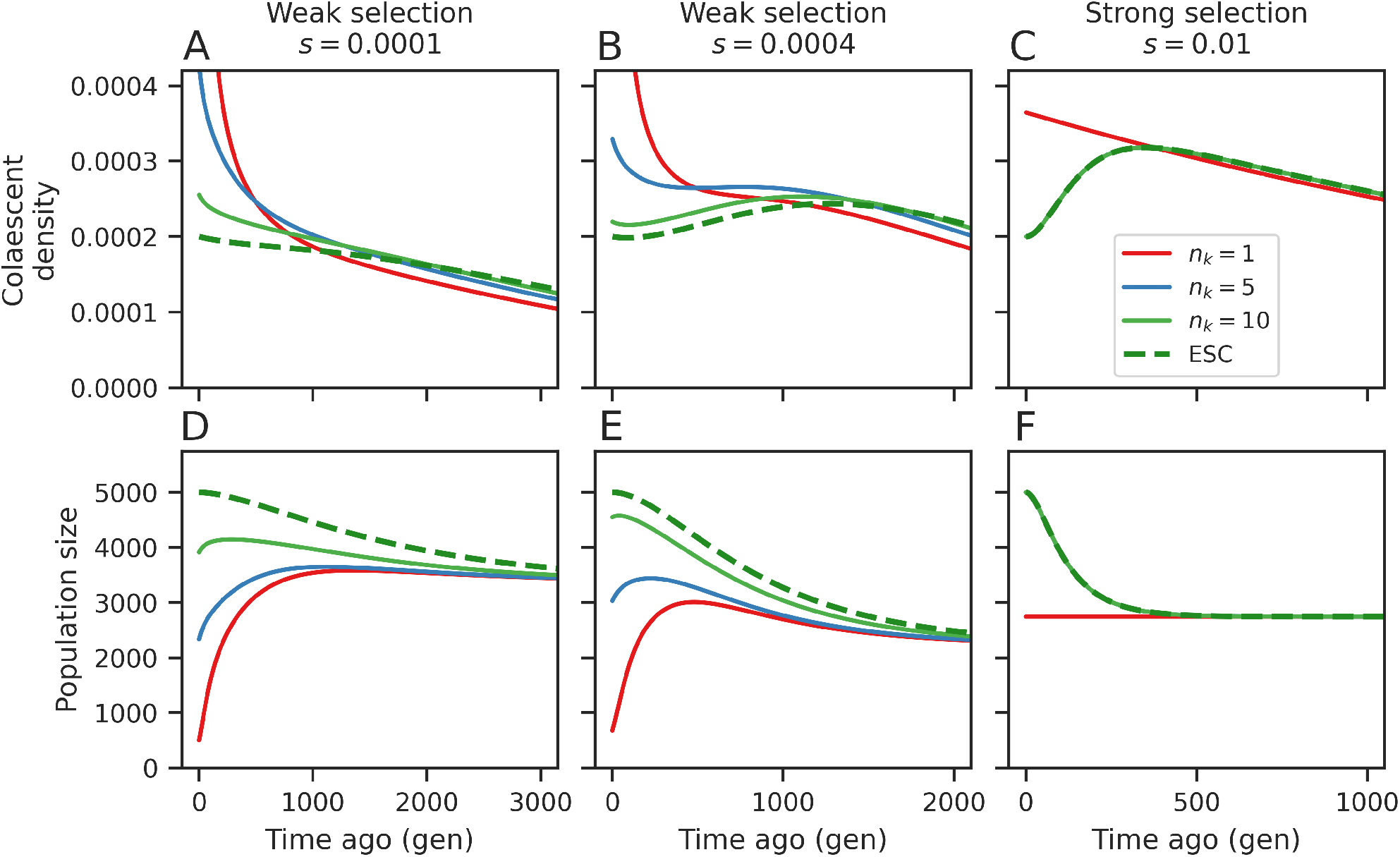
Expected coalescent times and population sizes under incomplete sampling. Predictions were calculated using the extended structured coalescent for *U*_*d*_ = 0.006, *N* = 5000 and three different selection coefficients (columns) when sampling from the *n*_*k*_ most frequent classes of the distribution of deleterious mutations (indicated colors) or the complete initial distribution (ESC). The selection coefficient *s* is indicated in the top panel headers.

In genomic regions without recombination, the chromosome genealogy represents a single realization of evolution. We posit that inference based on such samples may not be robust. Thus, in the next step, we explored the differences in estimates using two randomly sampled lineages from the mutation-load distribution. Samples of size two represent an extreme case of ascertainment, highlighting the challenges associated with small sample sizes. With this sampling scheme, we investigated the qualitative and quantitative effects of sampling lineages specific to particular mutation-load classes. Each sampled chromosome pair represents the expected genealogy from samples of two particular mutation-load classes. We calculated the probabilities of coalescence over time for these two lineages. We show the estimated history of effective population size for all possible pairs of samples (Figure S6). The inferred trajectories were arbitrarily categorized as corresponding to six simple types of demographic events (Figure S6). For example, we refer to a bottleneck if the population experienced a historical period of reduced size (Figure S6A, B, C). As previously described (Walczak *et al*. 2012), the estimated past effective population size histories are strongly dependent on the numbers of deleterious mutations in the classes from which the lineages are sampled. When lineages are sampled from different but closely neighboring fitness classes the estimated history of effective population size was categorized as a bottleneck, as bottleneck-like, or as an expansion (Figure 6, Fig. S6A-E, L-N). Our observations indicate a more pronounced bias toward bottleneck and bottleneck-like inferred demographies under conditions of weaker purifying selection (Figure S7) and when lineages are sampled from neighboring fitness classes (Figure S7).

**Figure 6.**
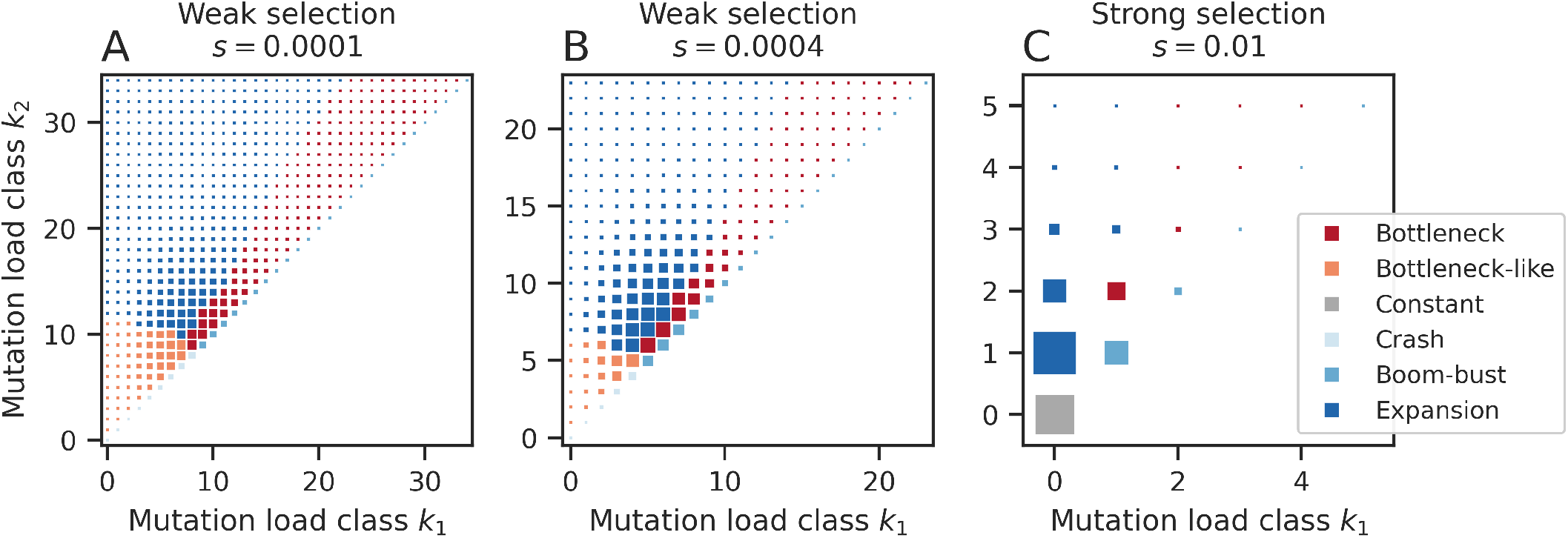
Expected demographic scenarios under the extended structured coalescent for samples of size two. Each data point represents an inferred demographic scenario (colors) of a pair of lineages sampled from the mutation-load distribution (x-and y-axis). The categories are shown for three different selection strengths *s* = 1 10^−4^ (A), *s* = 4 10^−4^ (B) and *s* = 1 10^−2^ (C) for *N* = 5000, *U*_*d*_ = 0.006. The size of the data points represents the probability of sampling a particular pair of mutation-load classes. Notably, the true demography is a constant population size.

In summary, when a population evolves under purifying selection, sampling schemes that inadequately represent the mutation-load distribution of the population may often result in qualitatively-biased inferences, such as transient increases or bottlenecks. These effects appear more pronounced for weaker purifying selection.

## Discussion

This study bridges an important gap in our knowledge of the coalescent process under purifying selection in non-recombining regions, previously limited to very weak or strong selection regimes. By extending the structured coalescent model, we incorporate weak purifying selection allowing deleterious mutations to occasionally fix in the population. This modification allows us to model the entire spectrum of selection strengths as a smooth transition between neutrality and strong selection, converging to the structured coalescent approach for strong purifying selection (Walczak *et al*. 2012; Nicolaisen and Desai 2012). The central idea of our approach is to consider relative mutation-load rather than absolute numbers of deleterious mutations, which allows us to incorporate occasional fixations by modifying migration rates in the structured coalescent.

Our extended structured coalescent reveals that the effective population sizes and coalescent distributions are intermediate between those observed under neutral and strong selection scenarios, but they show delayed equilibration under weaker selection (Figure 3, 4). Additionally, our study highlights how poor sampling strategy can lead to a misrepresentation of demographic events such as population expansions or crashes. This emphasizes the importance of considering purifying selection in demographic models, as well as the significance of representative sampling, particularly in the absence of recombination (Figure 5, 6). Our comprehensive analysis significantly enhances our understanding of evolutionary dynamics under varied selective pressures in non-recombining genomic regions and how these dynamics affect demographic inference.

In the model of Nicolaisen and Desai (2012), the mutation-load distribution at mutation-selection balance is stationary. That allows the time-reversal of the forward-in-time Markov process and, hence, a detailed description of the coalescent process. In our extended structured coalescent, this distribution is time-dependent due to the occasional fixation of deleterious mutations, and we therefore focus on the relative mutation-load distribution instead of the absolute numbers of deleterious mutations (Figure 2B). The rate at which deleterious mutations are fixed is the second key parameter in our approach. Together, these quantities allow us to describe the evolution of the ancestral lineage distribution backward in time. A key assumption of the structured coalescent model of Nicolaisen and Desai (2012) is that the ancestries of sampled lineages are independent. This assumption is justified under strong selection, as lineages move quickly to the zero mutation class where they mainly coalesce. In our extension of their approach, even when weak selection compromises the independence of lineages, this has minimal impact on the accuracy of our predictions (Figure 3, 4). Our results remain valid even when selection coefficients approach zero, and our model describes the mutation distribution of a neutrally evolving population (Figures 3A, 4A). In addition, the higher mean of the Poisson-like mutation-load distribution under weak selection is consistent with the lower selection efficacy in keeping deleterious mutations at low frequencies, particularly maintaining a high proportion of individuals in the least loaded class (Figure S3). Taken together, these findings suggest that using the mutation-load distribution and the rate at which mutations become fixed may suffice to describe the coalescent process in various selection scenarios.

Addressing the complexities of realistic distributions of fitness effects (DFE) poses a challenge for theoretical analyses. We therefor consider haploid populations and a single fixed selection coefficient to approximate the impact of deleterious mutations on fitness. We would argue that this is a reasonable approximation if the resulting mutation-load distribution under a more complex distribution of fitness effects can be accurately modeled using a single effective selection coefficient. For instance, Good *et al*. (2013) suggested that the effects of many weakly selected mutations on sequence diversity are identical to those of fewer strongly selected mutations and can be modeled collectively. We also expect that adding a proportion of strongly deleterious mutations will have little impact, as such mutations will be eliminated quickly and thus contribute little to the ancestry dynamics (Charlesworth *et al*. 1993). However, it is more difficult to predict how complex DFEs, where haplotypes carrying various mutations compete, may affect the coalescent process in non-recombining regions. The extent to which the mutation-load distribution and the fixation rate are sufficient to describe the coalescent process under more complex DFEs remains to be investigated in future studies.

Our results should readily extend to diploid populations with additive or multiplicative fitness effects. However, dominance and epistasis require a multi-layered structured coalescent that accounts for combinatorial mutation effects. A multi-layered structured coalescent model is unlikely to be amenable to a detailed mathematical analysis. Describing the mutation-load as a traveling wave in the continuous fitness space, where a mutation kernel reflects how lineages move through fitness space, appears as a promising but potentially challenging avenue for future research (Hallatschek 2011; Rouzine *et al*. 2003; O’Fallon *et al*. 2010).

Previous work on strong selection and recombination suggests that recombination does not affect the qualitative patterns of genealogy distortion and effective population size dynamics obtained in models without recombination (Nicolaisen *et al*. 2013). The recent growth of the effective population size due to purifying selection has also been shown in realistic simulations using various DFEs in diploid, recombining populations (Johri *et al*. 2021). An intriguing explanation for this observation is that a genome can be conceptualized as several independent non-recombining blocks (Weissman and Hallatschek 2013; Nicolaisen *et al*. 2013). We expect that our results extend to recombining organisms depending on the specific recombination rate in the investigated scenarios. In conclusion, the genealogy distortion and the history of effective population sizes generally fall between neutrality and strong selection for a broad range parameter combinations (Figure 3, 4).

Our results justify using methods that assume neutral evolution to infer the history of the effective population size, rather than the census size, in the presence of background selection. However, we highlight the potential ascertainment bias introduced by sampling schemes (Figure 5, 6). This bias is due to uneven sampling from the mutation-load distribution, such as neglecting rare haplotypes (Figure 5). The bias becomes more pronounced as purifying selection weakens. Describing the genealogies as the expected history of effective population sizes provides an incomplete picture as it comprises complex scenarios, ranging from expansion to recent crashes and bottlenecks. Thus, for ascertained sampling schemes, our extended structured coalescent model may help to capture the effects of purifying selection that might otherwise be mistaken for demographic events in the inferred effective population size history.

Our findings have significant implications for understanding the dynamics of coalescent history in scenarios with low or no recombination Genetic diversity must be considered within the context of specific chromosome models. For instance, purifying selection could partially explain the disparities in inferred demographic histories from uniparental markers in humans. Therefore, our study not only sheds light on the complexities of coalescent history and genetic diversity in non-recombining regions but also suggests that previous interpretations of human genetic diversity of uni-parental markers may need to be revisited to obtain a more nuanced understanding of human demographic histories.

## Significance statement

We developed an extended structured coalescent that accurately models weak purifying selection. We also discussed the limitations of using non-recombining genetic regions, such as uniparental markers in humans, for inferring past demography. Our study emphasizes the importance of accounting for the effects of purifying selection on neutral genetic diversity in non-recombining regions and avoiding sampling biases to achieve unbiased demographic inferences.

## Data availability

The simulation data from the analysis and the corresponding scripts are provided in https://github.com/sstruett/purifying_selection.git. The scripts to create the figures are provided in the same repository.

## Acknowledgments

The authors appreciate the CMPG members for their helpful comments and Dr. Kimberly Gilbert for her feedback on the manuscript.

## Funding

This work is funded by a Swiss NSF grant to Prof. Laurent Excoffier (No. 310030_188883), and the Interfaculty Bioinformatics Unit at the University of Bern, Switzerland.

## Conflicts of interest

The authors have no conflict of interest to declare.

## Supplementary Figures

**Supplementary Figure 1.**
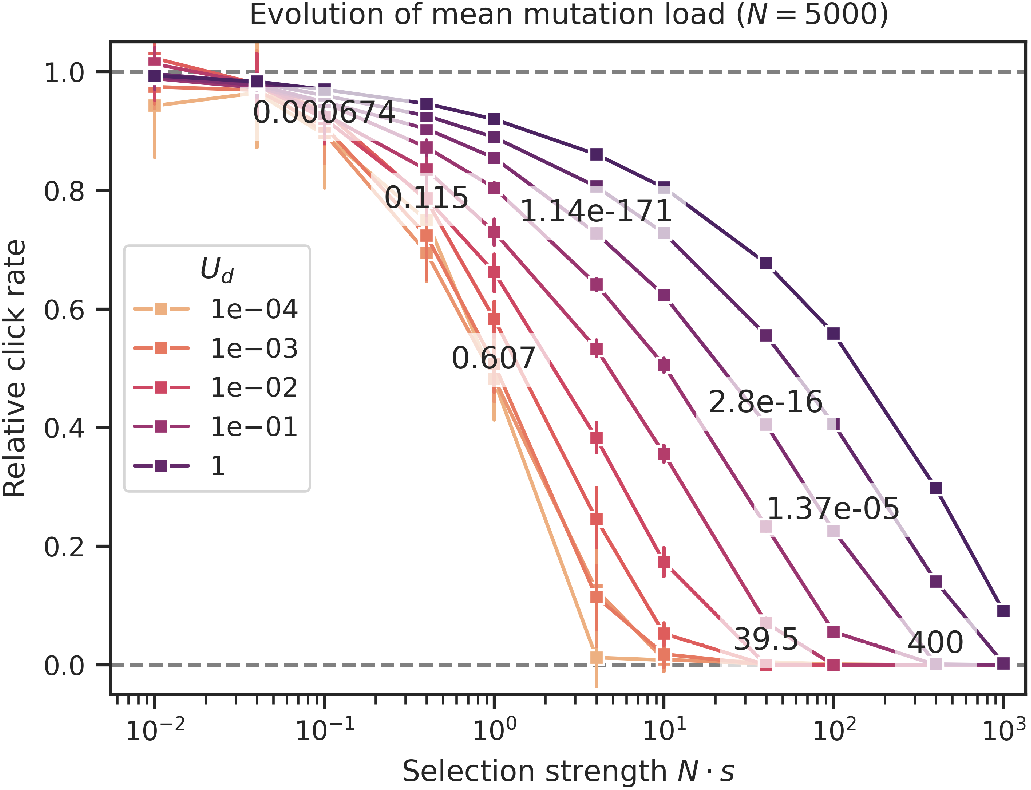
Click rates of populations under different parameters of *U*_*d*_ and *s* **for** *N* = 5000. We simulated the relative click rates, i.e., normalized to the deleterious mutation rate, of the evolving mutation-load distribution in populations under different parameters *U*_*d*_ and *s*. The error bars indicate the 95% confidence interval of the simulated data. For weak selection strength (*N s* < 1), the relative click rate approximates 1; in contrast, for stronger selection (*N s* ≫ 1), it approximates zero, indicating a successful escape from Muller’s ratchet due to effective purging. Intermediate values of the relative click rates under weak selection follow a smooth transition, where the exact shape depends on the indicated parameters. The numbers reported in the graph show *ϕ* values, indicating their negative correlation with the click rate.

**Supplementary Figure 2.**
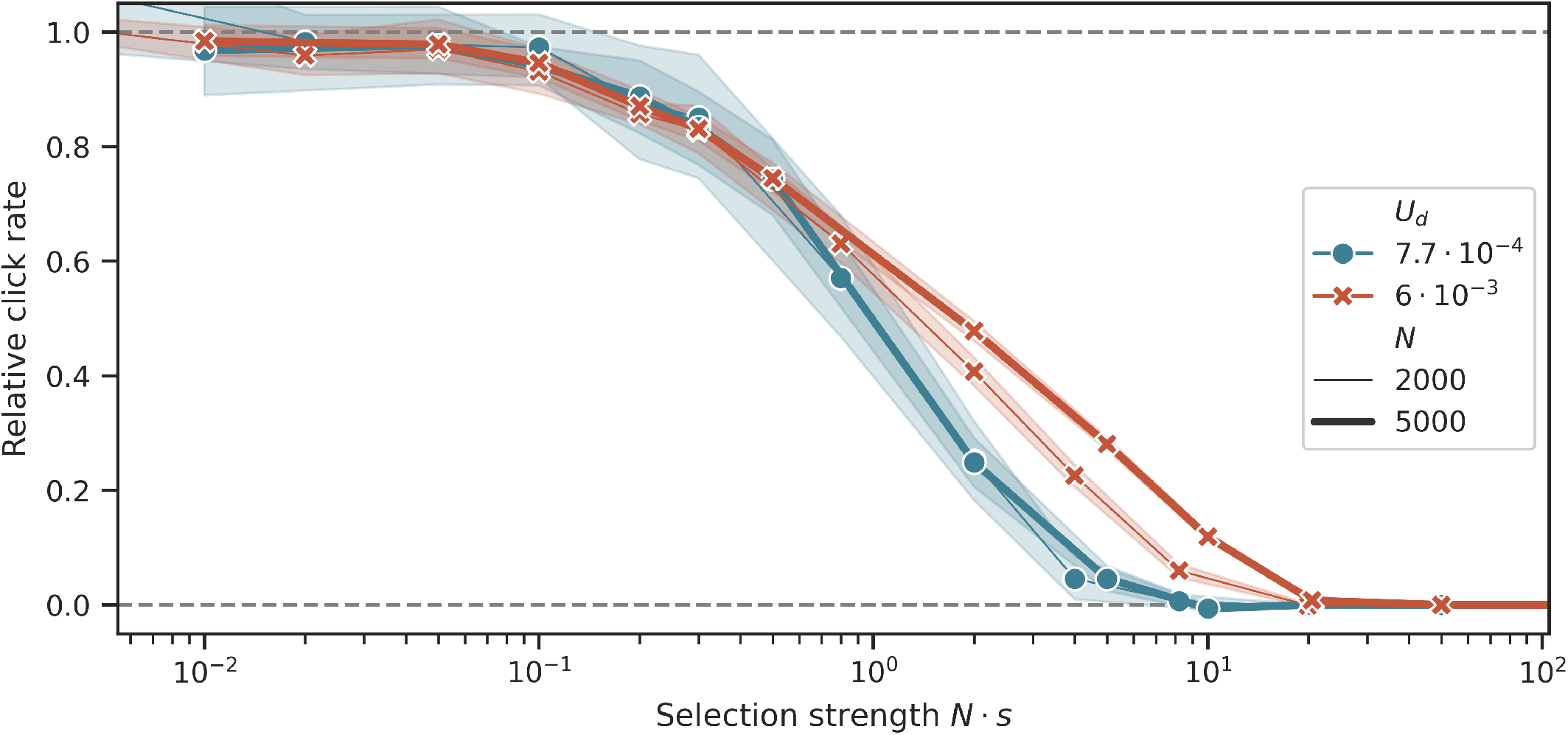
Simulated click rates of of populations evolving under purifying selection. The relative click rates, the click rate normalized to the deleterious mutation rate, of the mutation-load distribution under two different deleterious mutation rates *U*_*d*_ and two population sizes *N*_*e*_ depending on the selection strength. The bandwidth indicates the 95% confidence interval.

**Supplementary Figure 3.**
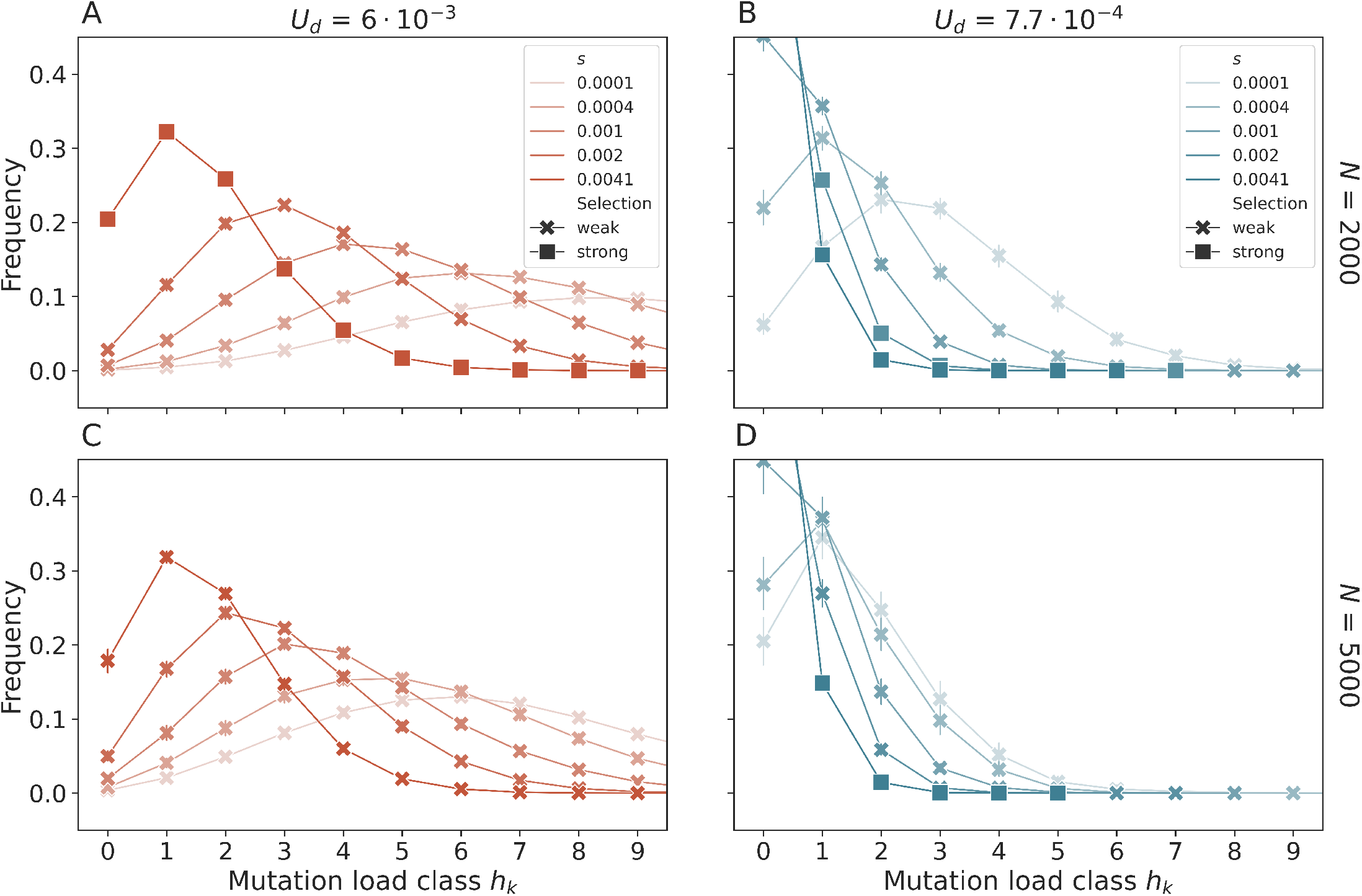
Simulated mutation-load distributions of populations evolving under purifying selection. The Poisson-like distribution of deleterious mutations is shifted toward higher mutation-load classes for either weaker selection or higher mutation rates. Scenarios of weak selection are those when the click rate *v* > 0. Strong selection scenarios correspond to a click rate of *v* = 0. The error bars indicate the 95% confidence interval of the simulated data. However, the variance is generally small, such that they may appear invisible.

**Supplementary Figure 4.**
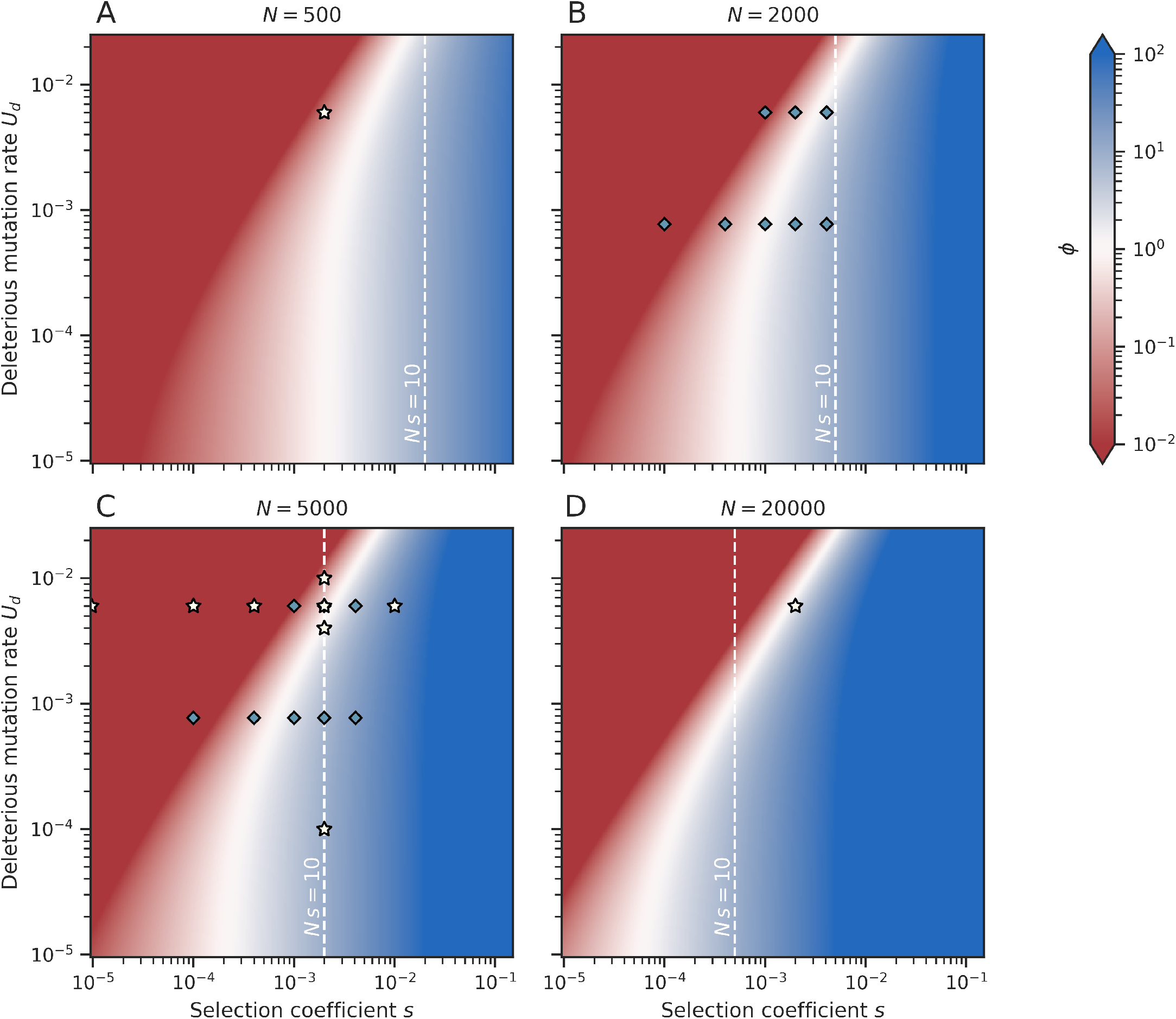
*ϕ* as a measure of the validity of the fitness class coalescent model. *ϕ >* 1 (white to blue) suggests strong selection and indicates that the assumptions of the structured coalescent model are valid. Conversely, values of *ϕ* < 1 (white to red) imply the presence of weak selection. The white dashed line represents the selection strength *N s* = 10. The diamonds and stars non-exhaustively indicate the simulated parameters from Figures 2, S1,S2,S3 (diamonds) and Figures 3, 4, S5 (stars).

**Supplementary Figure 5.**
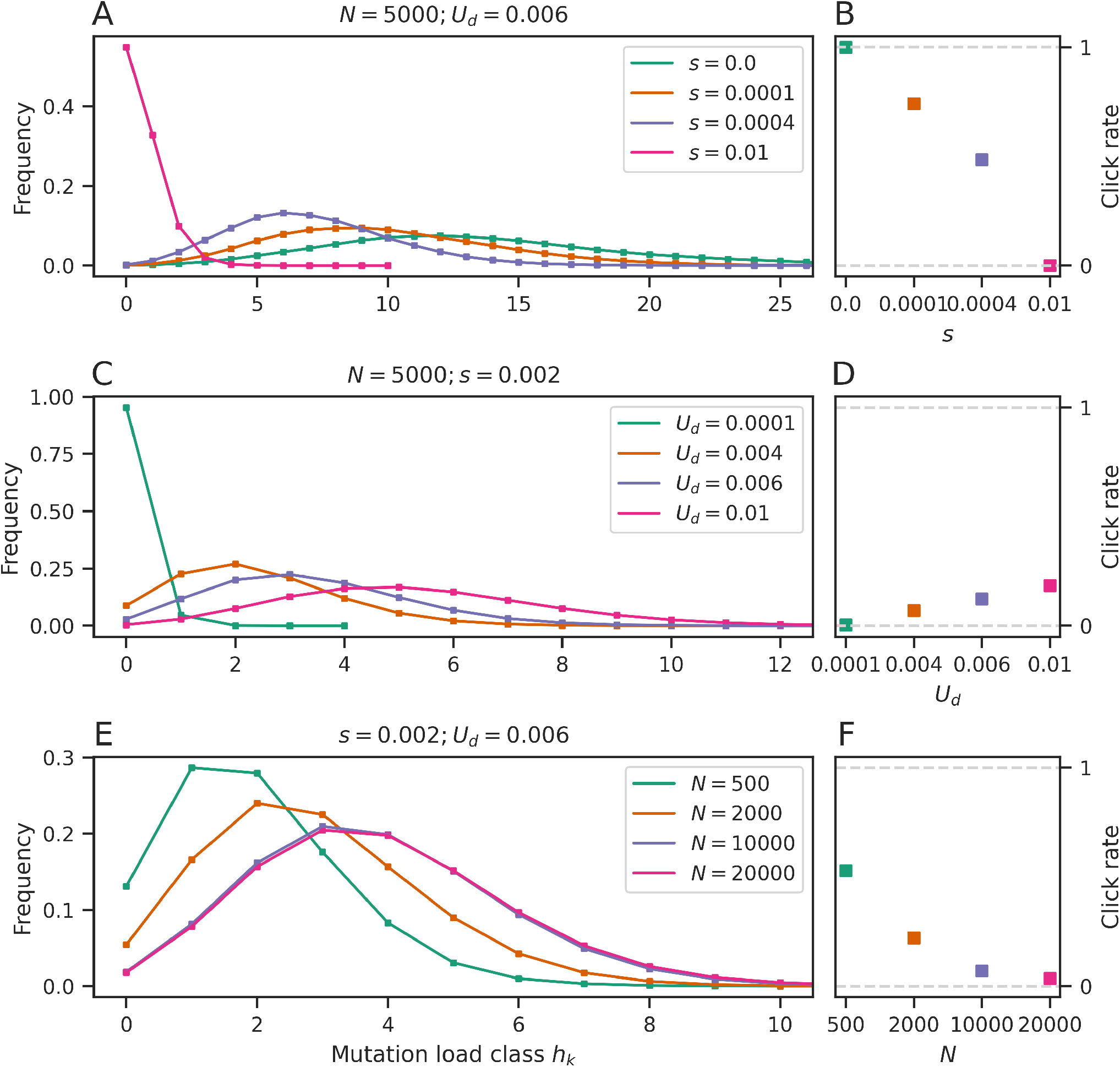
mutation-load distributions and click rates for the scenarios of Figure 3. This figure complements the coalescent densities in Figure 3 and provides the corresponding distribution of deleterious mutations and click rates for the same sets of parameters as indicated in the figure panels. Lower *s*, higher *U*_*d*_, and higher *N* parameter values can shift the mutation-load distribution toward higher mutation-load, which corresponds to larger click rates.

**Supplementary Figure 6.**
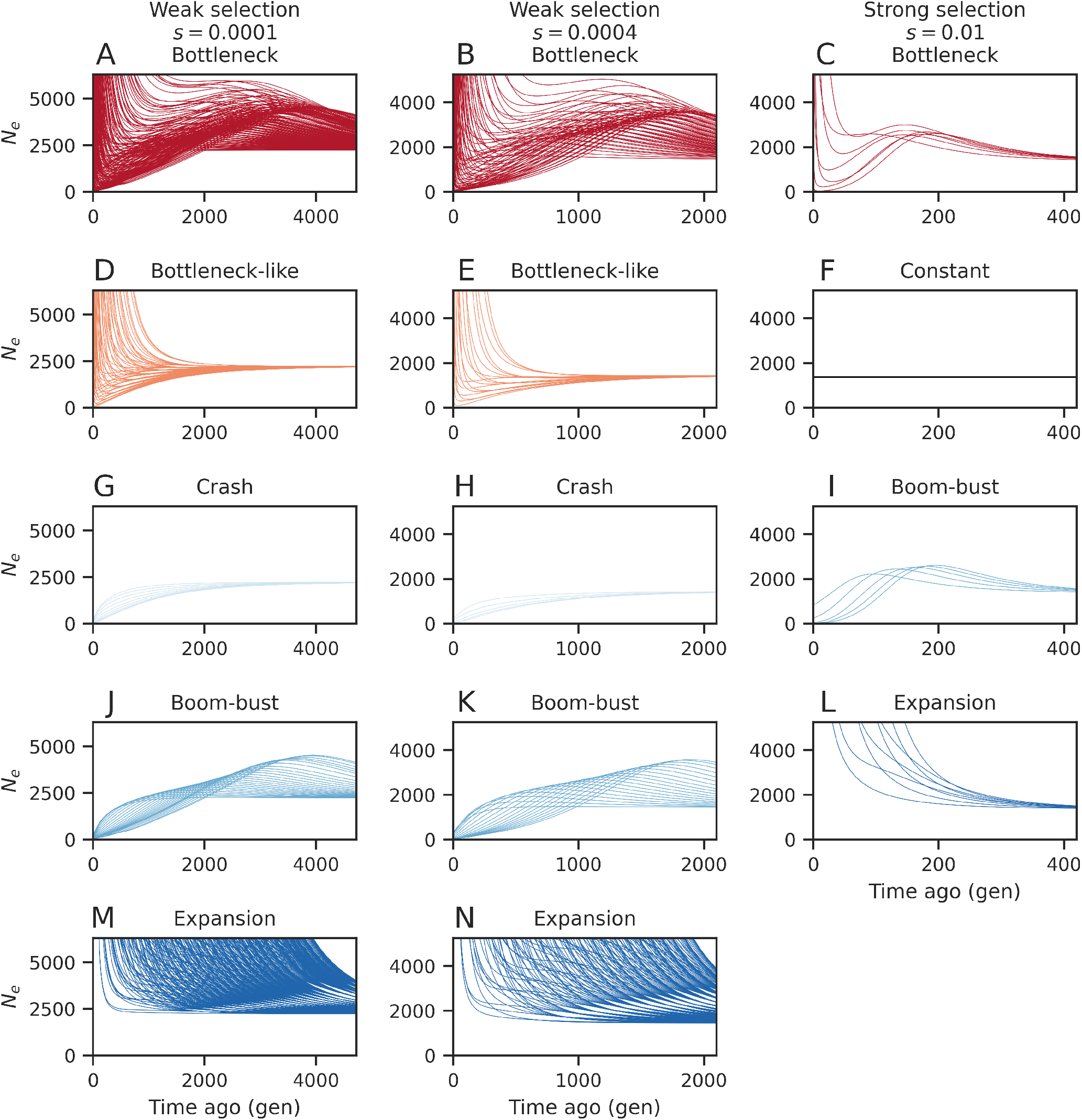
Expected demographic scenarios under the extended structured coalescent for samples of size two. Each line represents the expected history of effective population size of two randomly sampled lineages from the mutation-load distribution under the extended structured coalescent. The expected demographic scenarios are shown for *N* = 5000, *U*_*d*_ = 0.006 for three different selection strengths *s* = 1 10^−4^ (first column), *s* = 4 10^−4^ (second column) and *s* = 1 10^−2^ (third column). The inferred trajectories were arbitrarily categorized as corresponding to simple types of demographic events (see panel titles, colors).

**Supplementary Figure 7.**
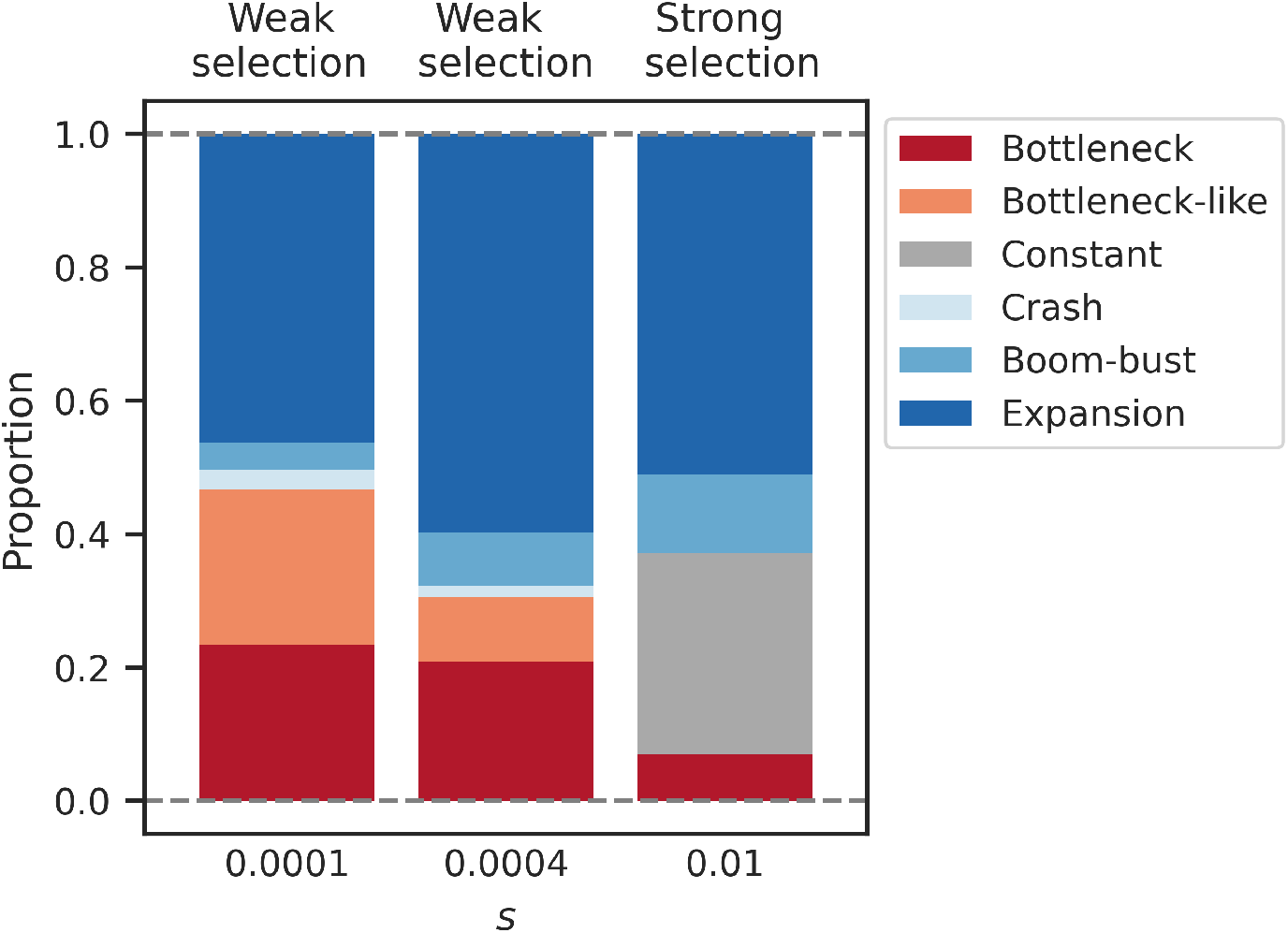
Distribution of expected demographic scenarios under the extended structured coalescent for samples of size two. The results of Figure S6 have been summarized to show the proportion of expected demographic categories for three different selection coefficients (x-axis) and *N* = 5000, *U*_*d*_ = 0.006.

## Notes

https://github.com/sstruett/purifying_selection.git

### Competing Interest Statement

The authors have declared no competing interest.

